# Intact reinforcement learning in healthy ageing

**DOI:** 10.1101/2023.05.25.542104

**Authors:** Wei-Hsiang Lin, Aaron M. Clarke, Karin S. Pilz, Michael H. Herzog, Marina Kunchulia

## Abstract

What does age in ageing? Results in reinforcement learning (RL) are mixed. Some studies found deteriorated performance in older participants compared to younger controls whereas other studies did not. Daniel et al. (2020) suggested that task demand can explain these differences, with less demanding tasks showing no effect of age. Here, we increased the task demand of previous studies turning them into a classic navigation task. We extracted 4 behavioral parameters and 2 parameters (learning and exploration rates) of a classic Q-learning model. Except for one specific parameter, all other parameters showed no group differences, i.e., RL turned out to be intact in older individuals also with higher task demands. It is important to publish such null results to avoid the stigmatizing impression of an overall performance deficit among older people.

## Introduction

To understand the mechanisms of ageing, it is of great importance to understand first what functions decline and which do not. Learning is often thought to significantly decay with age (Anguera et al., 2011; Salthouse, 2009). However, evidence is mixed. In reinforcement learning (RL), for example, a clear decline was found by Daniel et al. (2020) and van de Vijver & Ligneul (2020), whereas other studies found intact RL even though paradigms were rather similar (Eppinger et al., 2008; Lighthall et al., 2018). In these paradigms, an image is presented and participants push one of two buttons to receive a positive, negative, or neutral reward. Then, the next image is presented randomly from a set of given images, and so on. The task in these studies is not very demanding, provoking only light deficits in both the young and old group. For this reason, Daniel et al. (2020) have proposed that deficits of older people are only found in demanding tasks.

Here, we used a more demanding paradigm, where the presentation of image *n* was not chosen randomly, as in the above studies, but depended on the choice made by the participants at image *n-1*. One image was a goal image, and participants were asked to find it as often as possible within a certain period of time (Fig. 1). Hence, the experiment mimics a navigation task, which is more realistic than the above paradigms and involves additional aspects such as a systematic exploration of the RL environment, linking states not only to actions but also to other states. To increase task demand, we tested not only a short (0.5s) but also a longer (6s) inter-stimulus intervals (ISI) between images. Various parameters were extracted from the participants’ performance and a Q-learning model was fitted. To preface our results, with one exception, older participants’ performance was comparable to that of younger controls.

**Figure 1.**
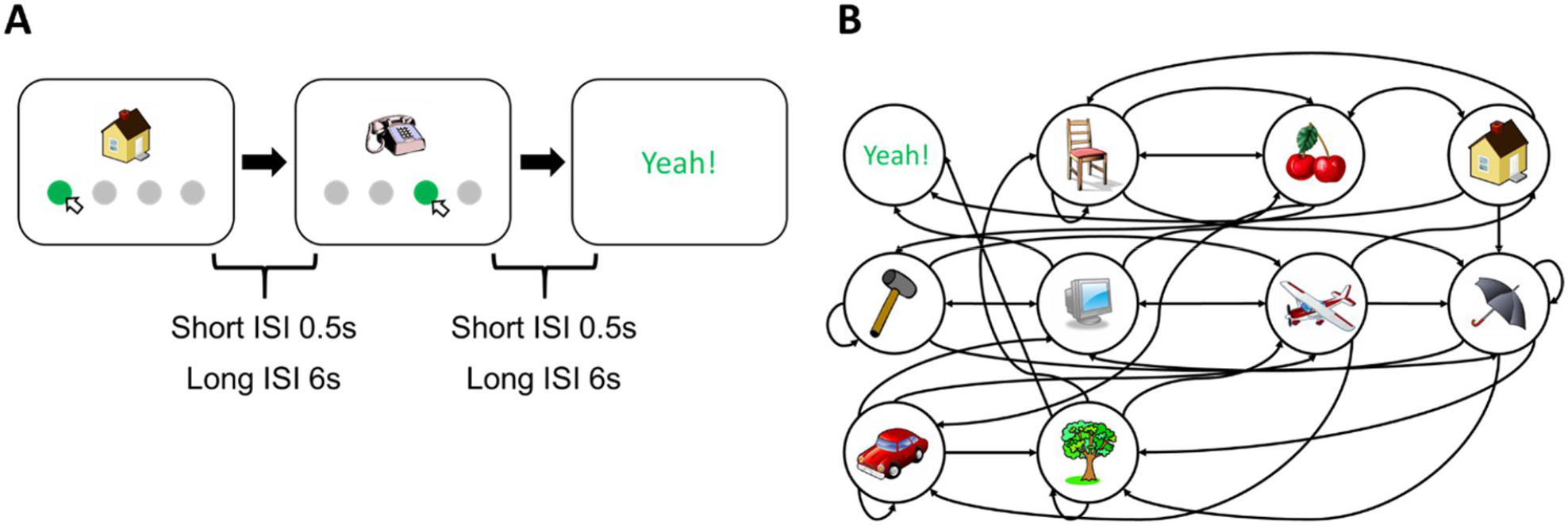
RL task. (A) An image was presented to the participants. Participants chose one of the disks below the image to proceed to the next image (state) until they found the goal state (“Yeah!”). Each participant performed under two different Inter-stimulus intervals (ISI) conditions, the short (0.5s) and long (6s). In experiment 1, participants had to reach the goal state as many times as possible within a limited duration (8 minutes for the short ISI and 30 minutes for the long ISI condition). In experiment 2, they had to find as many as possible goal states within 150 actions. Details can be found in the Methods and Materials. (B) Structure of the RL environment. Each image represents a state, and the direction of the arrow indicates the connection between different states. Importantly, the structure of the environment is very irregular in the sense that observers may go directly from image A to image B but not necessarily back. In all of the experiments, there are a total of nine states plus one goal state. As primary measures, we extracted 6 parameters including the number of episodes completed, the proportion of optimal actions, the improvement in both the number of episodes completed and the proportion of optimal actions, the learning rate and exploration rate (detailed in the Methods and Materials).

## Results

### Experiment 1

#### Effects of ISI and age

First, we examined whether performance was affected by age. Surprisingly, the number of episodes completed (Fig. 2A) and the proportion of optimal actions (Fig. 2B) did not differ significantly between young and old adults. Next, we divided all states into two categories, the adjacent and distant states, based on their proximity to the goal state (Fig. S1A). Again, there was no significant main effect of age neither in the accuracy of the adjacent states (Fig. S1B, left) nor the distant states (Fig. S1B, right). Hence, older adults performed equally well as young adults.

**Figure 2.**
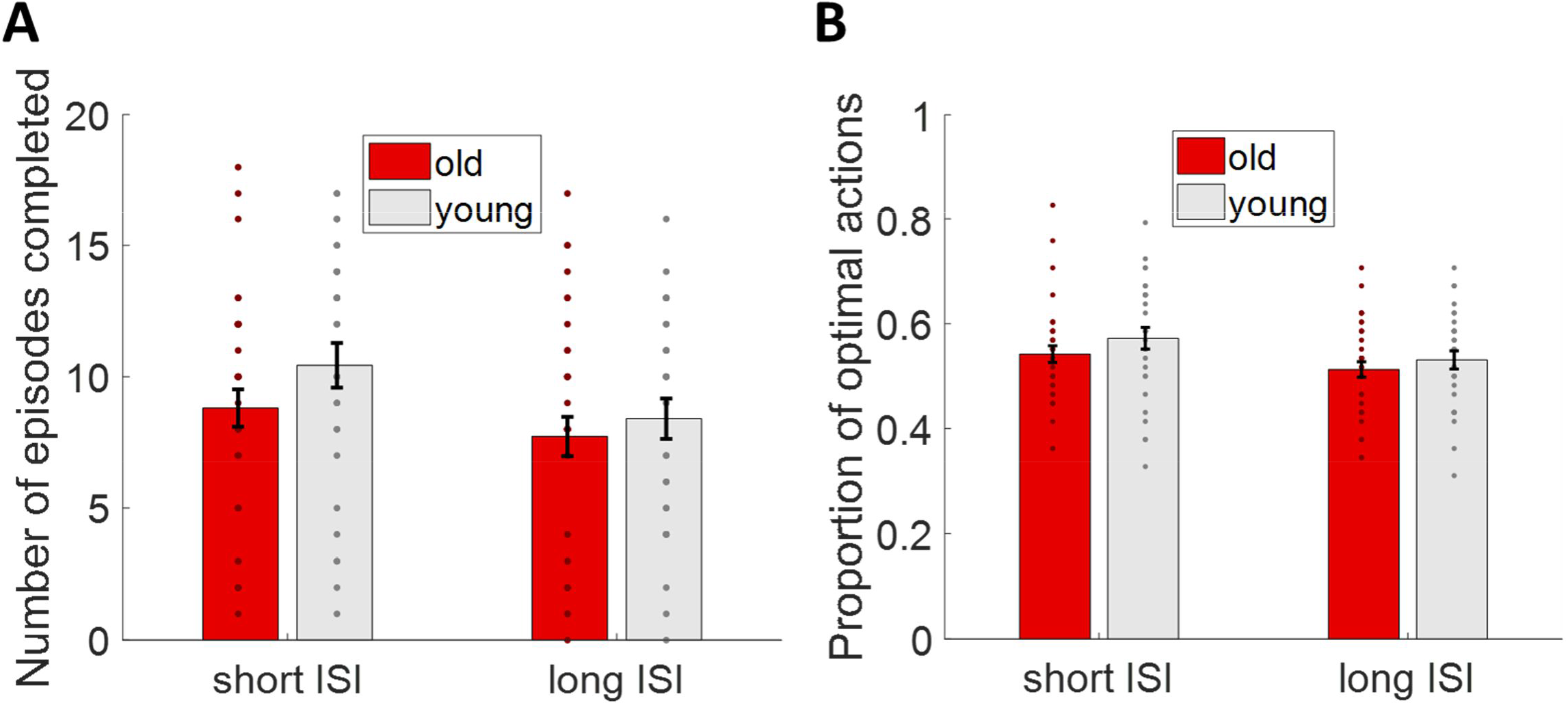
Performance in the RL task. (A) The number of episodes completed in each ISI condition for each age group. Dots indicate the performance of individual participants. (B) The proportion of optimal actions in each ISI condition for each age group. Error bars represent ±1 SEM.

#### Improvement of performance

We next investigated the improvement of performance during the RL task by dividing the trials into two sets, the first 29 trials and the last 29 trials of each observer. There was a significant interaction between the two groups and ISI when it came to the number of episodes completed. A post-hoc test showed that the long ISI condition drove the interaction (Fig. 3A). A slight trend but no statistically significant improvement in performance was observed (Fig. 3B). For a closer examination, we fitted the accuracy data across trials with a log function which yielded two parameters, the slope and intercept. Both the intercept and the slope did not differ between the groups (Fig. S2A).

**Figure 3.**
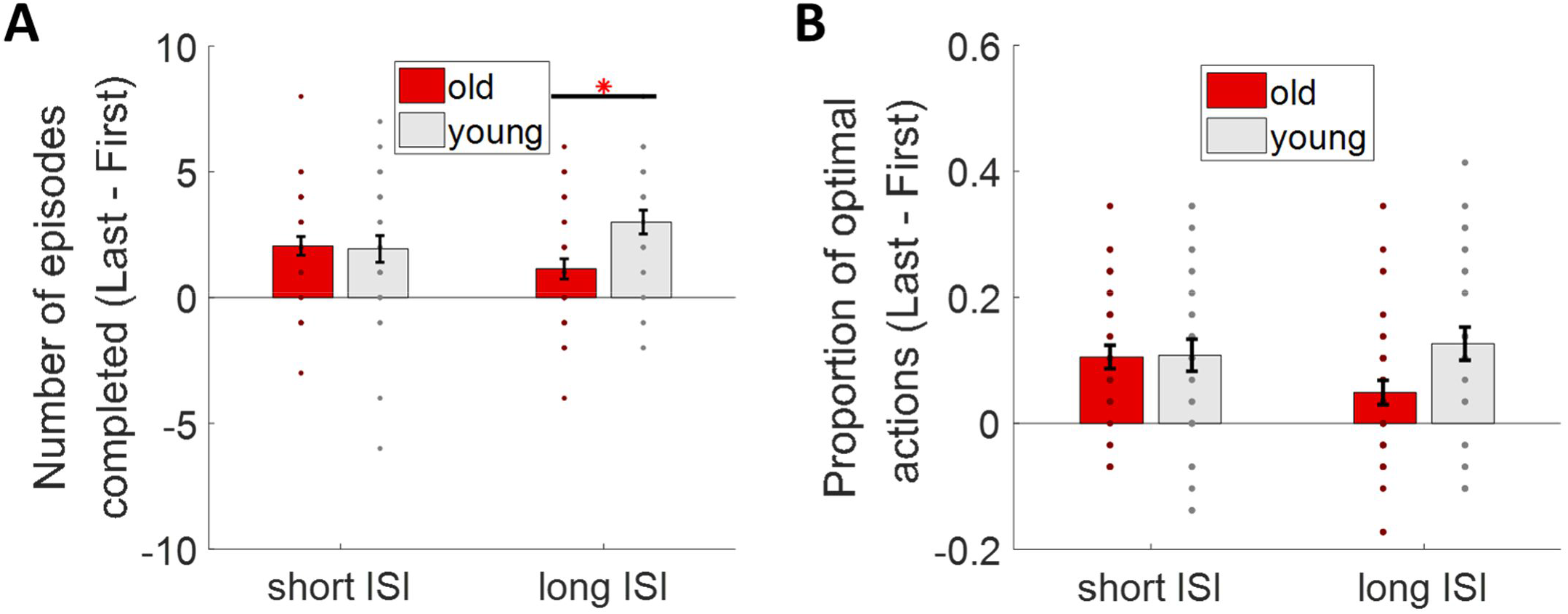
Improvement of performance. The improvement in the accuracy for each age group and ISI condition. (A) The improvement in number of episodes. *: p < 0.05 by post-hoc Tukey’s test. (B) Improvement in proportion of optimal actions. Error bars represent ±1 SEM.

#### Perseveration behavior and action entropy

We assessed the extent of suboptimal action selection by quantifying the action entropy, which is a measure of the randomness of actions. Older adults had a lower improvement in action entropy compared to young adults (Fig. S2B). We suggest below that the difference may be caused by memory deficits.

Additionally, we also examined the perseveration behavior, which is characterized by a persistent repetition of suboptimal actions. There were no significant differences between young and older adults (Fig. S2C, left; Fig. S2C, middle). Furthermore, the improvement in perseveration behavior was not significant across the age groups (Fig. S2C, right).

Our findings indicate a difference in the improvement of action entropy between age groups, with the older group showing less improvement compared to the younger group, regardless of the ISI condition.

#### Q-learning

Both learning rates (Fig. 4A, top) and the inverse temperature (Fig. 4A, bottom) did not show significant differences between the two age groups. The learning rate reflects the speed of learning, and the inverse temperature reflects the randomness of choices or exploration rate. The results suggest that both groups revealed similar learning efficiency and exploration through the RL environment.

**Figure 4.**
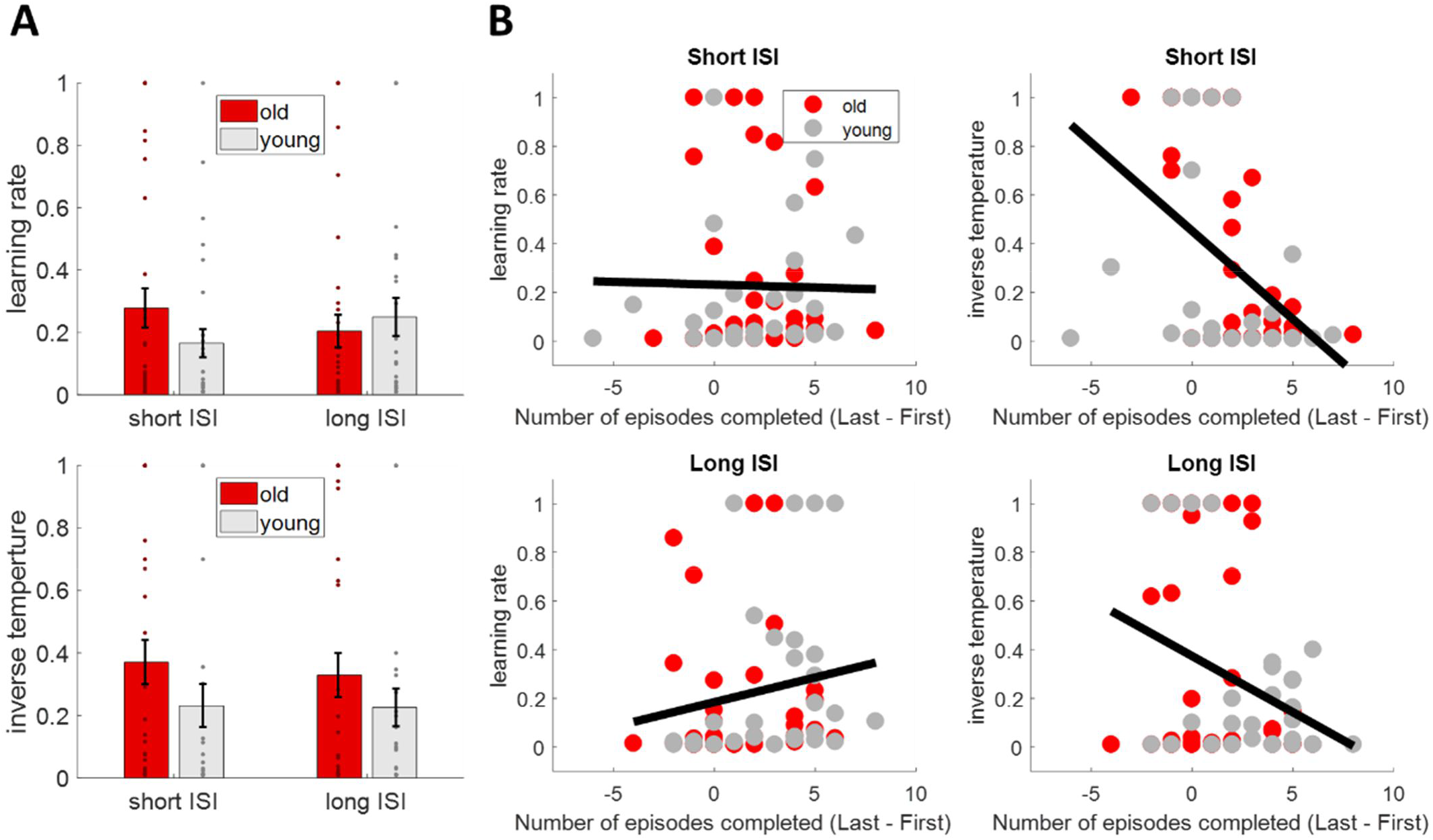
Q-learning model. The two parameters retrieved from the Q-learning model for each condition and for each age group: (A) Top: The learning rate; Bottom: The inverse temperature. The conventions are the same as in Fig. 2. (B) Correlation between the parameters and the improvement in performance. Top panel: The correlation between the improvement in performance and the learning rate (left) and the inverse temperature (right) in the short ISI condition. Bottom panel: The correlation between the improvement in performance and the learning rate (left) and the inverse temperature (right) in the long ISI condition. The black line is the linear regression fit of the parameters and performance improvement combined for both older and young adults.

We next linked performance to the Q-learning model to study whether there are correlations between the model parameters and the improvement in performance. Interestingly, we found that only the learning rate in the long ISI condition positively correlated with improvement in performance (Fig. 4B, Bottom left). The inverse temperature was negatively correlated with improvement in performance merely in the short ISI condition (Fig. 4B, Top right), though the same negative trend was also present in the long ISI condition (Fig. 4B, Bottom right).

Effect size and statistical values can be found in Table S1.

### Experiment 2

#### Effects of ISI and age

Contrary to experiment 1, a significant interaction of ISI and age was observed in the proportion of optimal actions (Fig. 5A, right), but a marginal difference in the number of episodes completed (Fig. 5A, left).

**Figure 5.**
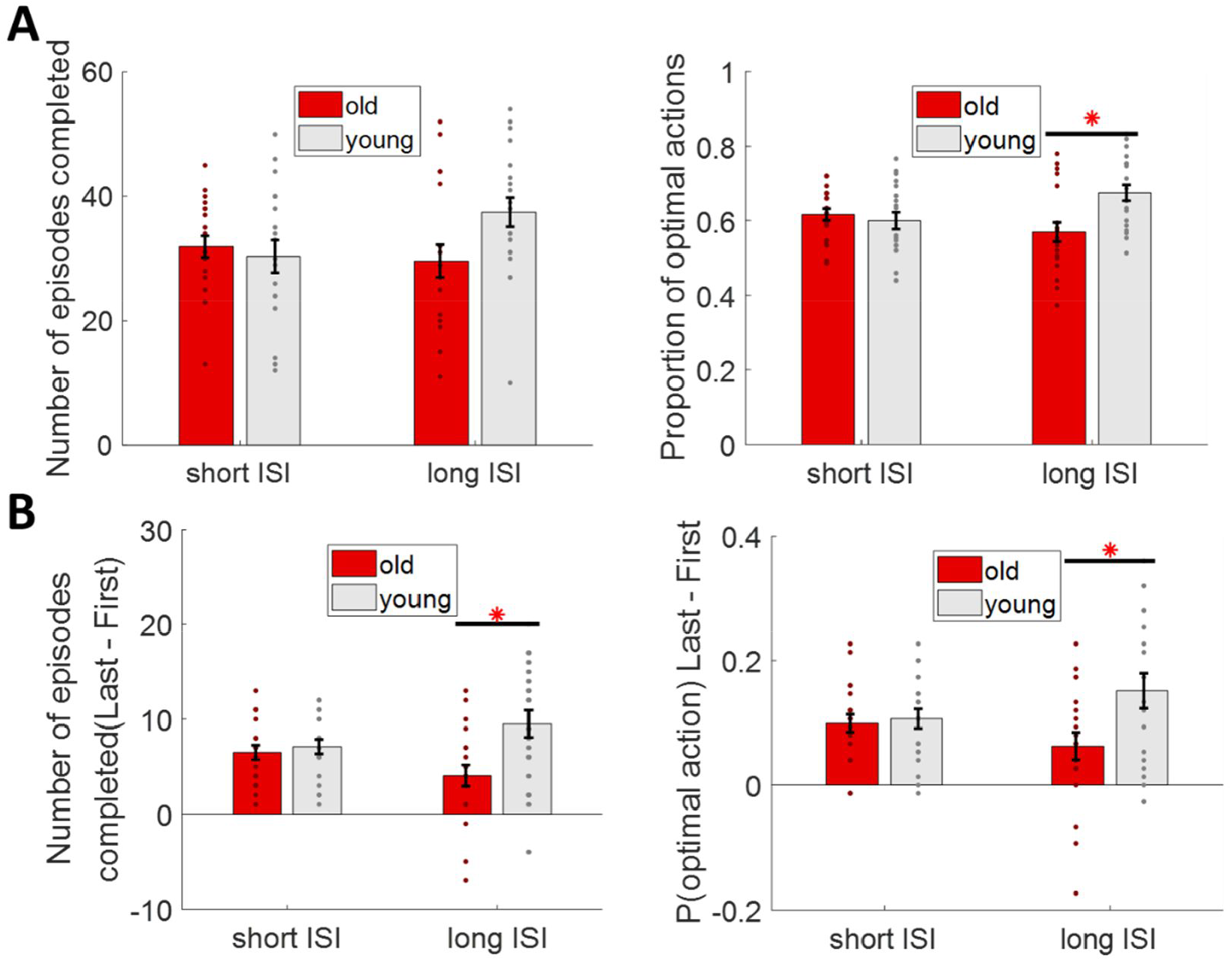
Performance in experiment 2. The conventions are the same as Fig. 1. Error bars represent ±1 SEM. (A) Left: The number of episodes completed in each ISI condition for each age group. Right: The proportion of optimal action in each ISI condition for each age group. *: p < 0.05 post-hoc Tukey’s test. (B) The improvement in the accuracy for each age group and ISI condition. Left: The improvement in number of episodes. Right: Improvement in proportion of optimal actions. *: p < 0.05 by post-hoc Tukey’s test.

Moreover, there was a significant difference in improvement in performance in the long ISI between older and young adults (Fig. 5B). For adjacent states, there were no significant difference in accuracy was observed in the adjacent states (Fig. S3A, top), but a significant difference appeared in the distant states (Fig. S3A, bottom).

This is further supported by an improvement in action entropy: the young group improved more than the old only in the distant states (Fig. S3B, top). These observations are consistent with the pattern in experiment 1. Additionally, the improvement in action entropy is highly correlated with the improvement in performance (Fig. S3B, bottom) in the long ISI condition, indicating that the lack of improvement in action entropy in the old group is influencing performance.

The results indicate that only in the long ISI condition older and younger adults performed differently. The difference in accuracy can be explained by the fact that the older participants were less accurate in the distant states.

#### Questionnaire

The results of the questionnaire revealed that both older and young adults were able to accurately recognize the images used in the RL task, with no participants making mistakes. Next, we measured the confidence rating of their answers. The confidence ratings were based on a scale where a rating of 1 represented high confidence and a rating of 4 indicated high uncertainty. Interestingly, there was no significant difference in the confidence ratings between older and young adults in the adjacent states (Fig. S3C, top). However, a clear main effect of age in the distant states (Fig. S3C, bottom).

#### Q-learning

Similar to the results of experiment 1, both learning rates (Fig. 6A, top) and the inverse temperature (Fig. 6A, bottom) did not differ between the two age groups. This suggests that, while there may be some subtle differences in the learning processes of older and younger adults, overall there is not significant difference in the parameters used for Q-learning. However, despite the absence of group differences, the inverse temperature was found to be significantly correlated with accuracy (Fig. 6B, Top right) rather than learning rate (Fig. 6B, Top left). This indicates that while the learning rate may not have a direct effect on performance, the inverse temperature, which controls the exploration-exploitation trade-off, does affect the accuracy.

**Figure 6.**
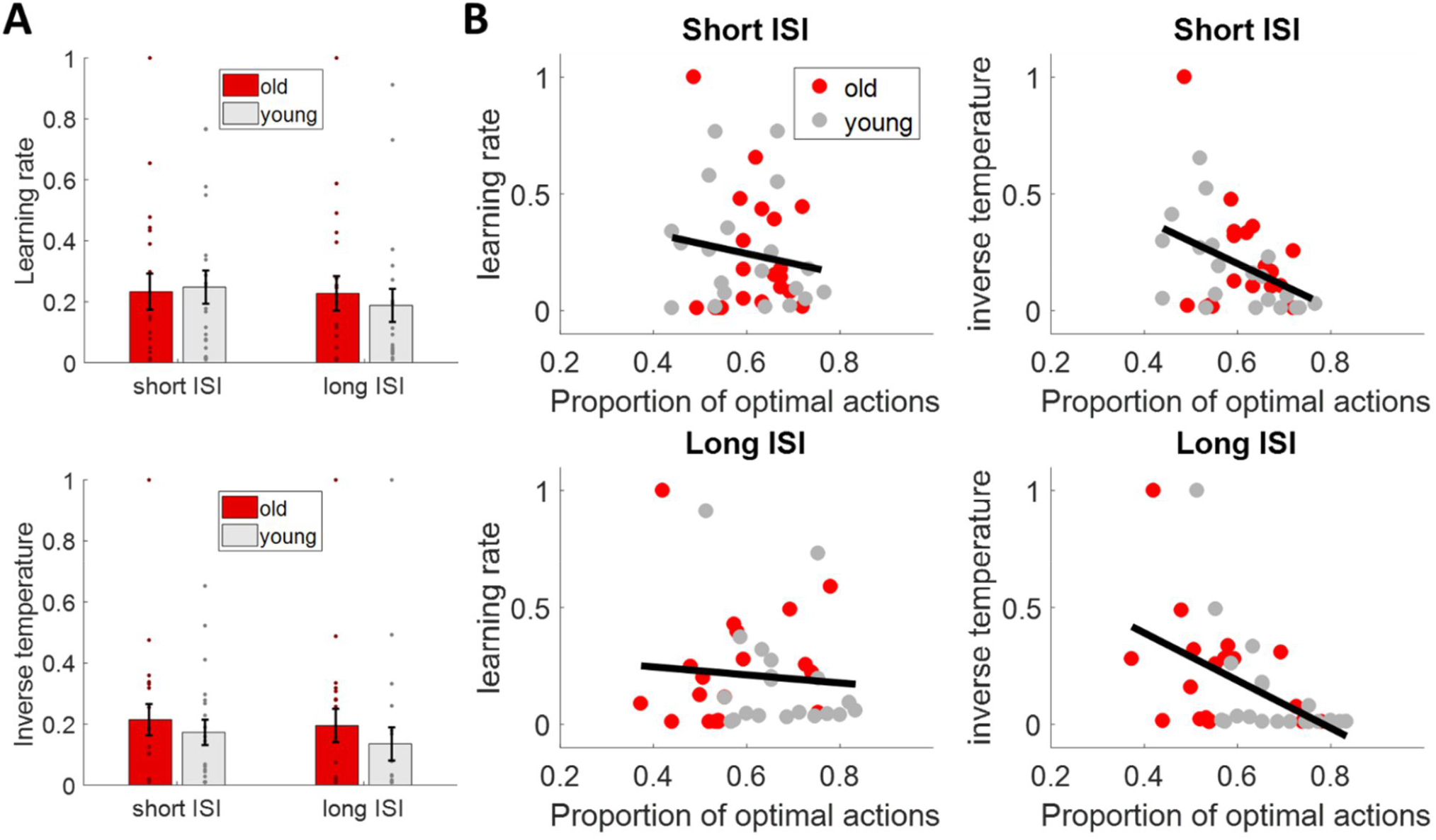
Q-learning model and the performance – experiment 2. The conventions are the same as in Fig. 4. A Q-learning model was fitted to the data and two parameters retrieved from the model for each condition and for each age group: (A) Top: The learning rate; Bottom: The inverse temperature. (B) The correlation between the parameters and the proportion of optimal actions. Top panel: The correlation between the proportion of optimal actions and the learning rate (left) and the inverse temperature (right) in the short ISI condition. Bottom panel: The correlation between the proportion of optimal actions and the learning rate (left) and the inverse temperature (right) in the long ISI condition.

Effect size and statistical values can be found in Table S2.

## Discussion

In the current study, we aimed to investigate whether older adults exhibit deficits in RL. In experiment 1, older adults performed as well as young adults with the number of episodes completed, proportion of optimal actions, improvement in proportion of optimal actions, learning rate and the exploration rate. In experiment 2, we found significant differences in the long ISI condition for improvement in accuracy and the secondary measures derived from this measure. Overall, the results of our study are consistent with previous studies (Lighthall et al., 2018; Pietschmann et al., 2011) but not others (Daniel et al., 2020). Hence, RL is largely intact in older observers with our paradigm.

Moderate to large effects were observed in the measurements reflecting the performance differences between the two age groups in the long ISI condition. There are several explanations: a genuine RL deficit, a working memory deficit, or stronger fatigue. First, none of the parameters of the Q-learning model was abnormal in the older participants, including learning and exploration rates, which speaks rather against an RL deficit.

Second, in the memory questionnaire, older adults demonstrated less confidence in recognizing distant states. In addition, the improvement of action entropy was less pronounced. These findings may speak indeed to a slight memory deficit. Lighthall et al. (2018), employed a RL task also with short and long ISIs. Although the authors did not observe behavioral differences between the two age groups for either ISI condition, they found a significant difference in the hippocampal activity pattern between the age groups in the long ISI condition. Potentially the higher memory load in the long ISI condition leads to differences in hippocampal activation, but does not manifest on the behavioral level because of the simpler task.

Third, it is well known that older people fatigue more quickly than younger ones (Enoka & Duchateau, 2016). Hence, instead of memory load, higher fatigue in the older population may explain the results in the long ISI condition. However, our analysis of adjacent and distant states shows that only the distant states exhibited differences in terms of the proportion of optimal actions, indicating that the older adults were still able to locate the correct actions when the goal state was close enough. Hence, a memory deficit seems to be the best explanation at the moment.

As mentioned in the introduction, Daniel et al. (2020) claimed that older people perform worse than younger ones in demanding but not simpler tasks. The had an easy and a harder condition and found no performance differences in the easy condition but a trend for a group comparison in the hard condition. However, the conclusion is potentially not valid because of a ceiling effect in the easy condition (performance between younger and older controls: 97% vs. 94%). Potentially, there is a group effect also in the easy condition but definitely in the harder condition. Since task demand seems not to be the crucial aspect, there must be other reasons for the differences in the paradigms.

One of the limitations in the ageing field is that the older population exhibits a high variance due to differences in ageing. In addition, the mean age is often different. Small sample sizes may lead naturally to sampling biases, which may lead to different outcomes. In addition, there are demoscopical, socio-economical, and cultural differences. Hence, it may simply be the case that our and others null results come from a too “healthy” and/or still too “young” population (our mean age was 68 and 66 years and in Daniel et al. (2020) it was 70 years). Thus, not the paradigms and differences between paradigms are key but sampling of the population.

Next to performance, learning and exploration rate, we also tested for perseverations, a behavioral pattern where individuals continue to perform repeated action sequences regardless of whether they are optimal or not. We did not find an increased perseveration rate in older adults, contrary to previous results, for example, in the Wisconsin Card Sorting test (Shaqiri et al., 2019). With the very same paradigm used in this study, we tested schizophrenia patients and found evidence for perseverations compared to age-matched controls. Hence, our paradigm is sensitive to perseverations. Perseveration of schizophrenia patients are often attributed to abnormal dopamine levels leading to suboptimal action selection (Durstewitz & Seamans, 2008). In RL models, dopamine levels are classically related to the learning rate (Sutton & Barto, 2018) and prediction error (Schultz et al., 1997; Wise & Rompre, 1989). If this were all true, our results indicate an intact dopaminergic system.

In summary, our findings indicate that reinforcement learning can remain relatively preserved during ageing. This holds true for a large range of outcome measures we determined including learning and exploration rate in classic Q-learning models. We emphasize the importance to publish such null results. Suppressing null results may, otherwise, create the impression that older people are deficient in almost all paradigms, which does not seem to be true.

## Acknowledgements

This project was funded by the Swiss National Science Foundation (SNF) NCCR project Synapsy: The Synaptic Bases of Mental Diseases and the Sinergia project “Learning from Delayed and Sparse Feedback”.

## Materials and Methods

### Experiment 1

#### Participants

Forty older healthy adults and thirty healthy young adults were recruited from the Free University of Tbilisi, Georgia. Detailed demographical information is presented in Table 1.

**Table 1.** The demographic information of participants.

| Experiment | Experiment 1 |  | Experiment 2 |  |
| --- | --- | --- | --- | --- |
| Group | Young | Old | Young | Old |
| Age | 25.03±4.2 | 68.75±8.3 | 21.8±3.1 | 66.3±5.5 |
| Gender | M=14, F=16 | M=17, F=23 | M=9, F=11 | M=5, F=15 |
| Education years | 17.2±2.7 | 14.26±3.3 | 15.15±1.9 | 13.95±2.8 |
| MoCa | NA | 27.18±1.1 | NA | 27.75±1.5 |

#### Task

Participants performed a reinforcement learning task (Fig. 1). Participants were presented with one image in the center of the screen, accompanied by four gray disks below. Clicking on one of the disks brought the participants to the next image. We call the clicks sometimes “actions” and the images “states” in accordance with RL terminology. Participants had no time limit for choosing a disk. The objective was to find a goal image, labeled with the word “Yeah!”. Before the start of the experiment, all nine possible images were presented on the screen. The goal image was not presented but participants were informed about it. Observers initiated the experiment by clicking on a gray disk at the bottom of the screen.

We used two inter-stimulus intervals (ISI) of 0.5 seconds (short ISI condition) and 6 seconds (long ISI condition) between the participant’s responses and the next state, respectively. The objective of the task was to reach the goal state as frequently as possible within a limited time of 8 minutes for the short ISI condition, and 30 minutes for the long ISI condition. The order of the two ISI conditions was counterbalanced among the participants, i.e., half began with the short ISI condition and the half started with the long ISI condition. For each of the two ISI conditions, there were two distinct transition matrices, determining the transitions from one image to the next depending on the actions of the participants. The same matrices were used for all participants.

### Experiment 2

#### Participants

Twenty older healthy adults and twenty young healthy adults were recruited from the Free University of Tbilisi, Georgia. Individual who had previously taken part in the first experiment were no eligible. Detailed demographical information is presented in Table 1.

#### Task

Experiment 2 follows a similar procedure as experiment 1, with the difference that, instead of a fixed duration of 8 min, the number of trials was fixed. Accordingly, the objective of the task was to reach the goal state as many times as possible within the given number of trials. Furthermore, the order of the two ISI conditions was fixed, with the long ISI condition always coming after the short ISI condition.

Immediately after the RL task, a memory task was given to the participants. In total, there were eighteen images. Twelve of these images were the same as in the RL task, the other six images were novel. For each of the images, participants were asked 1) to indicate whether the given image appeared in the RL task, and 2) to rate their confidence in question one, on a 4-Likert scale.

### Behavioral analysis

#### Data pre-processing

In experiment 1, participants were instructed to reach the goal state as often as possible within 8min. Due to the nature of the task, different observers visited a different number of images ranging from 30 to 200 due to differences in decision-making and reaction times. To ensure the comparability of the data among the participants, only the first 58 trials (the fifth percentile of the total number of trials among all participants) were used. Any trials exceeding this threshold were discarded. Four participants from the older population were removed from the study, as their total number of trials was lower than 59 trials. It is important to note that this pre-processing procedure only applied to experiment 1 as in experiment 2 the number of trials was fixed for all participants.

#### Behavioral performance

We determined the number of episodes completed and the proportion of optimal actions taken. The number of episodes completed refers to the number of times the participants reached the goal state. The proportion of optimal actions is the number of times a participant chose the that optimally reduced the distance to the goal state from the current state divided by the total number of actions performed in the task. Furthermore, we assessed their improvement in performance by calculating the difference in the last and first 29 actions for the number of episodes completed and the proportion of optimal actions. A larger difference indicates better improvement, thereby implying enhanced learning. These four measures are our primary measures. Please note, these measures are not independent of each other. To gain further insights we used a number of secondary measures that are also not independent from the primary measures and can be seen as sub-measures.

First, the nine states of the environment were further categorized into adjacent and distant states. Adjacent states are states that were only one step away from the goal state. The six distant states were at least two steps away from the goal state. We analyzed performance separately for the two measures.

Second, to gain more insight into how performance progressed over time, we calculated the cumulative probability of optimal action across all trials. For each participant, we fitted a log function to estimate the intercept and the slope of the performance curve:

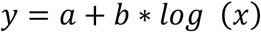

*x* represents trial numbers, *a* and *b* are the intercept and the slope, respectively, *y* is the cumulative proportion of optimal actions in each trial.

Third, we tested perseveration behavior. It is essential to efficiently select the optimal actions for a given state in order to achieve superior performance. Choosing repeatedly a non-optimal action in a given is suboptimal and may be related to perseverative behavior or memory deficits. We determined perseverations in two ways. First, we determined the average length of repeating action sequences. We averaged the length of these repetitions of actions across all episodes for each participant. For instance, an episode with the actions “1,2,3,1,2,3,4,2,1” has an average perseveration of two because the action sequence “1,2,3” appeared twice. Second, we calculated the proportion of repeated state-action pairs. In an episode, we extracted all the states and the corresponding actions to determine the probability that the same state-action pair has been visited within the same episode. In order to be considered optimal, a state should not be visited multiple times within an episode.

Fourth we determined the action entropy, measuring the randomness of the actions chosen. The theoretical max entropy is

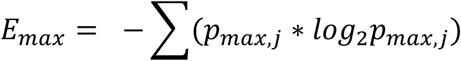

The probability distribution of completing each of the four actions was then determined for each state. The entropy of each state is

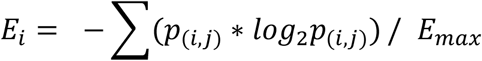

Here, *i* represents states, ranging from one to nine, while *j* represents actions, ranging from one to four. Averaging the action entropy across all states was computed for each participant. High action entropies indicate poor action choices (Sojitra et al., 2018).

#### Computational modelling

A Q-learning model (Sutton & Barto, 2018) was used to quantify learning:

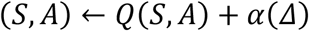

*S* represents the state of the current trial, *A* is the action taken in the current trial, *Δ* is the prediction error, *Q* is the Q-value for the given state and action, and *α* is the free parameter of the learning rate. Whenever a participant performed a new action, the Q value was updated by the prediction error, which is defined as the difference between the current reward and the expected reward:

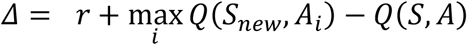

*r* stands for the reward, *S*_*New*_ is the new state after performing action *A* in the given state *S*. The probability of choosing an action is then determined by a soft-max rule:

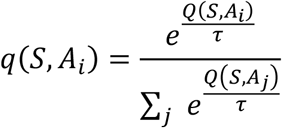

*q* is the probability of choosing action *A* given state *S*. τ is the free parameter normally referred to as inverse temperature which corresponds to the exploration rate.

#### Statistical analysis

We conducted a two-by-two repeated measures ANOVA with age group (old and young) and ISI (short and long) as independent variables. We quantified the relationship between measurements using Pearson correlations, except for the relationship between measurements and Q-learning model parameters, which were calculated with Spearman’s correlation due to the non-normal distribution of the parameters.

The primary parameters, focusing on performance, learning, and Q-learning model in RL, are presented in the main text. The secondary parameters, are found in the supplementary figures. All details are summarized in Table S1, Table S2, and Table S3.

## Supplementary figures

**Figure S1.**
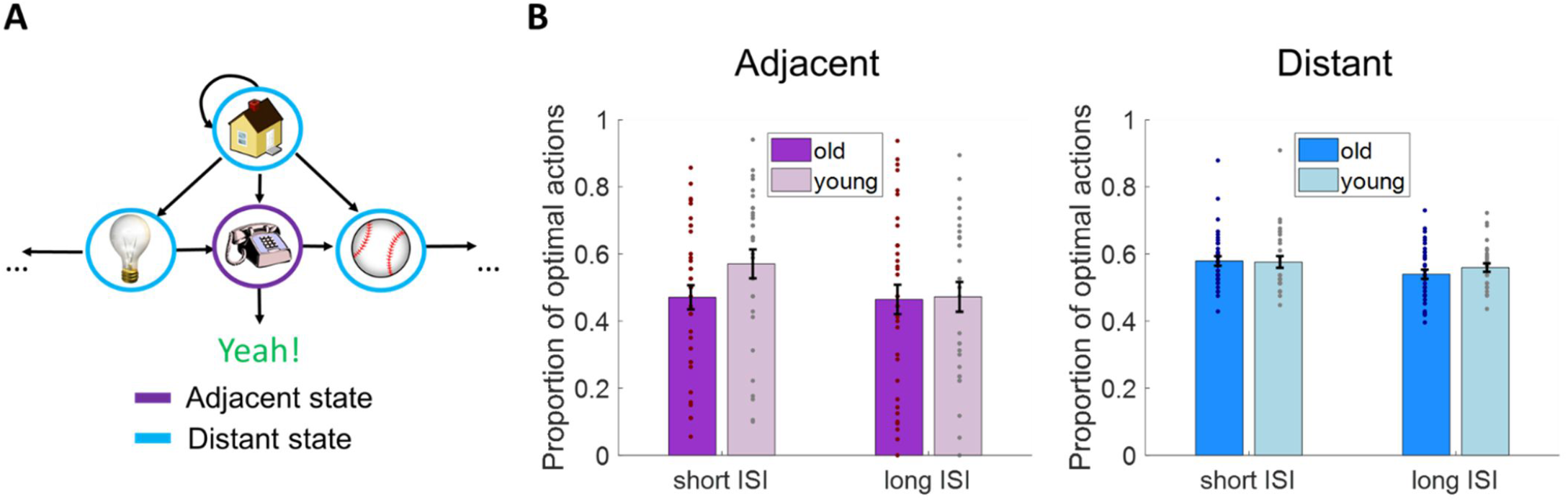
Performance in the RL task. (A) The adjacent state is indicated in purple, and refers to a state that is directly connected to the goal state. Distant states, indicated in blue, are those that require multiple steps to reach to the goal state. (B) Left: The proportion of optimal action for the adjacent states. Right: The proportion of optimal action for the distant states. Error bars represent ±1 SEM.

**Figure S2.**
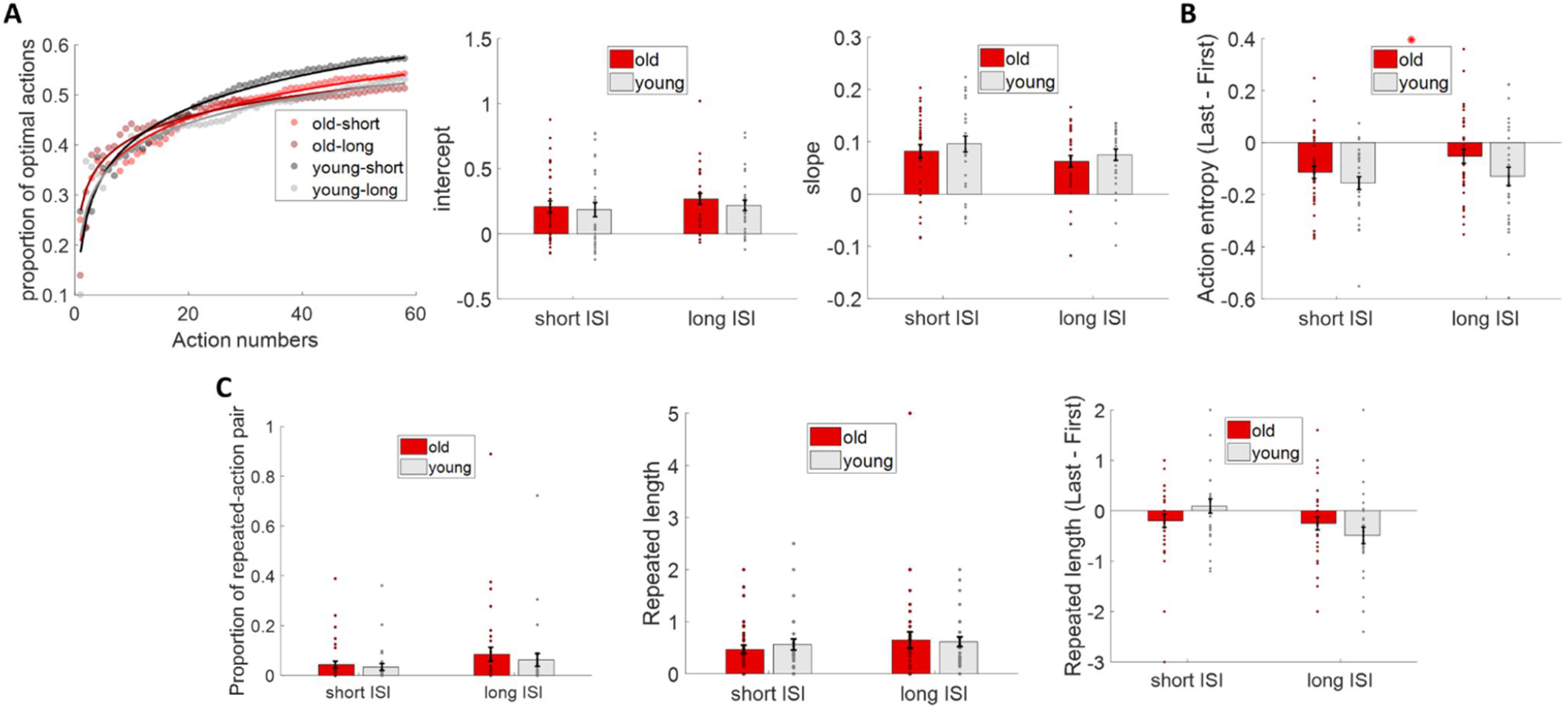
Perseveration behavior and action entropy. (A) Left: The cumulative proportion of optimal actions across time. The two ISI conditions and age groups were plotted separately. Each dot represents the average cumulative proportion of optimal actions across the participants in each age group. The solid line shows the fit of the log function. Right: The two bar graphs depict the slopes and intercepts of the fitted log function. (B) The improvement in action entropy. A main effect of age is presented. *: p < 0.05. (C) Left: The proportion of repeated state-action pair of each age group in each ISI condition. Middle: The repeated action length of each age group in each ISI condition. Right: The improvement in the repeated action length. Error bars represent ±1 SEM.

**Figure S3.**
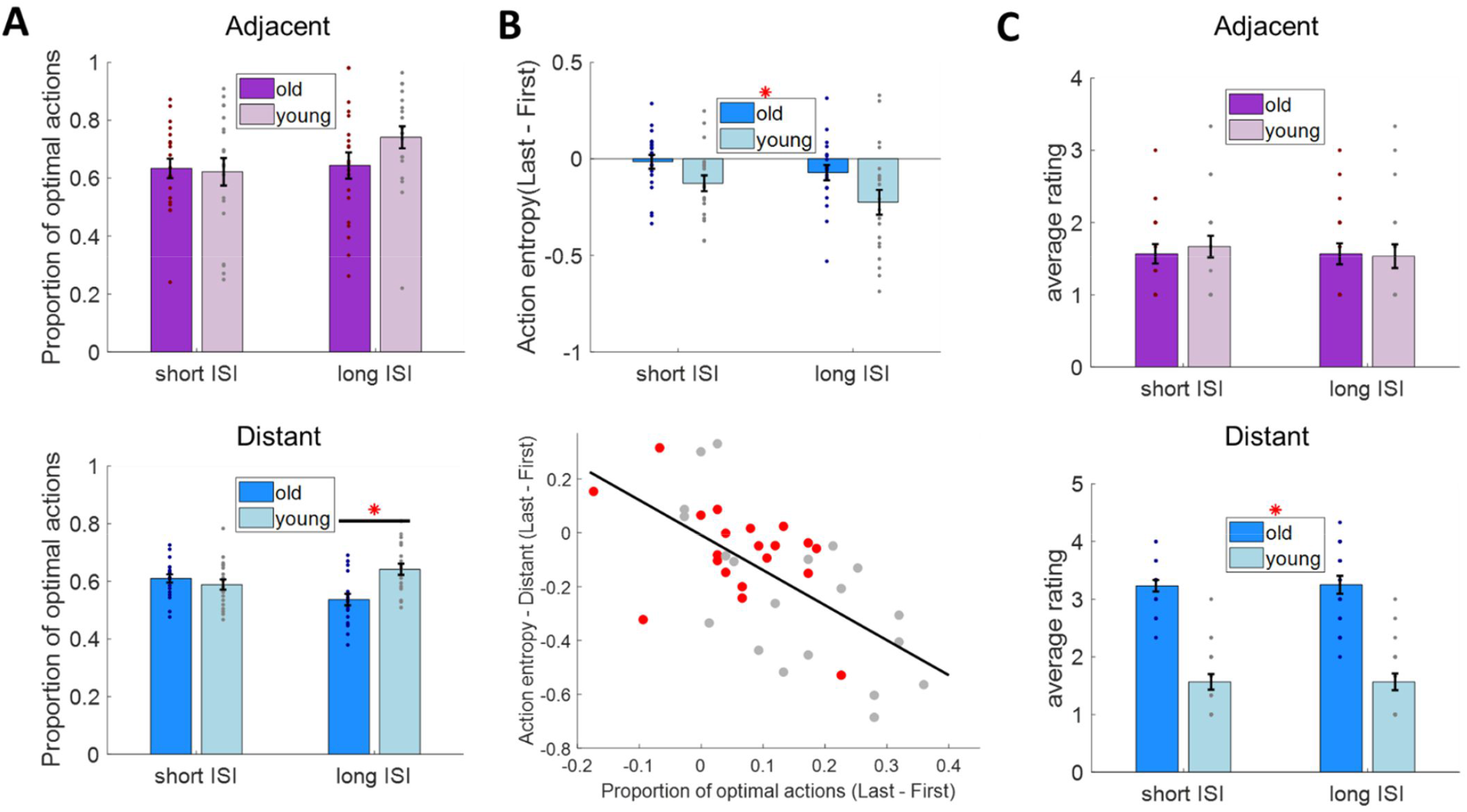
Secondary measurements for experiment 2. (A) Top: The proportion of optimal action in each ISI condition for each age group in the adjacent states. Bottom: The proportion of optimal action in each ISI condition for each age group in the distant states *: p < 0.05 post-hoc Tukey’s test. (B) Top: The improvement in action entropy in distant states. Bottom: The correlation between the improvement in performance and the improvement in action entropy in the distant states. (C) Confidence ratings of the questionnaire. Top: The average confidence ratings in adjacent states for each age group in each ISI condition. Bottom: The average confidence ratings in distant states for each age group in each ISI condition. Error bars represent ±1 SEM.

**Table S1.** Summary of parameters for experiment 1.

| Primary parameters – experiment 1 | Short ISI | Long ISI |
| --- | --- | --- |
| Number of episodes completed (2A) | age: $F(1,64) = 1.98$, $p = 0.16$, partial $\eta^2 = 0.03$<br>interaction: $F(1,64) = 0.44$, $p = 0.51$, partial $\eta^2 = 0.007$ | |
| Proportion of optimal actions (2B) | age: $F(1,64) = 1.83$, $p = 0.18$, partial $\eta^2 = 0.03$<br>interaction: $F(1,64) = 0.14$, $p = 0.71$, partial $\eta^2 = 0.002$ | |
| Improvement in number of episodes completed (3B) | Age: $F(1,64) = 3.66$, $p = 0.06$, partial $\eta^2 = 0.054$<br>interaction: $F(1,64) = 5.4$, **$p = 0.02$**, partial $\eta^2 = 0.078$ | |
| | ns | post-hoc test **$p_{\text{tukey}} = 0.018$**<br>Cohen's $d = 0.74$ |
| Improvement in proportion of optimal actions (3B) | age: $F(1,64) = 3$, $p = 0.09$, partial $\eta^2 = 0.045$<br>interaction: $F(1,64) = 3.1$, $p = 0.08$, partial $\eta^2 = 0.046$ | |
| Q-learning-alpha (4A) | age: $F(1,64) = 0.39$, $p = 0.54$, partial $\eta^2 = 0.006$<br>interaction: $F(1,64) = 1.77$, $p = 0.19$, partial $\eta^2 = 0.027$ | |
| Q-learning-tau (4A) | age: $F(1,64) = 2.98$, $p = 0.09$, partial $\eta^2 = 0.044$<br>interaction: $F(1,64) = 0.07$, $p = 0.79$, partial $\eta^2 = 0.001$ | |
| Secondary parameters – experiment 1 |  |  |
| Proportion of optimal actions – adjacent (S1B) | $F(1,64) = 1.48$, $p = 0.23$, partial $\eta^2 = 0.023$<br>interaction: $F(1,64) = 0.07$, $p = 0.25$, partial $\eta^2 = 0.02$ | |
| Proportion of optimal actions – distant (S1B) | age: $F(1,64) = 0.33$, $p = 0.57$, partial $\eta^2 = 0.005$<br>interaction: $F(1,64) = 0.63$, $p = 0.43$, partial $\eta^2 = 0.01$ | |
| Intercept of log fitting (S2A) | $F(1,64) = 0.65$, $p = 0.42$, partial $\eta^2 = 0.01$ | |
| Slope of log fitting (S2A) | $F(1,64) = 1.15$, $p = 0.29$, partial $\eta^2 = 0.02$ | |
| Improvement in action entropy (S2B) | age: $F(1,64) = 5.21$, **$p = 0.026$**, partial $\eta^2 = 0.075$;<br>interaction: $F(1,64) = 0.37$, $p = 0.55$, partial $\eta^2 = 0.006$ | |
| Proportion of repeated-action pair (S2C) | age: $F(1,64) = 0.06$, $p = 0.81$, partial $\eta^2 = 0.0009$<br>interaction: $F(1,64) = 0.39$, $p = 0.53$, partial $\eta^2 = 0.006$ | |
| Repeated action length (S2C) | age: $F(1,64) = 0.59$, $p = 0.45$, partial $\eta^2 = 0.009$<br>interaction: $F(1,64) = 0.09$, $p = 0.77$, partial $\eta^2 = 0.001$ | |

**Table S2.** Summary of parameters for experiment 2.

| Primary parameters – experiment 2 | Short ISI | Long ISI |
| --- | --- | --- |
| Number of episodes completed (5A) | age: $F(1,38) = 1.55$, $p = 0.22$, partial $\eta^2 = 0.04$<br>interaction: $F(1,38) = 4.84$, $p = \mathbf{0.03}$, partial $\eta^2 = 0.11$ | |
|  | ns | ns |
| Proportion of optimal actions (5A) | age: $F(1,38) = 3.55$, $p = 0.07$, partial $\eta^2 = 0.08$<br>interaction: $F(1,38) = 9.77$, $p = \mathbf{0.003}$, partial $\eta^2 = 0.2$ | |
| | ns | post-hoc test $p_{\text{tukey}} = \mathbf{0.005}$<br>Cohen's $d = -1.1$ |
| Improvement in number of episodes completed (5B) | age: $F(1,38) = 10.1$, $p = 0.003$, partial $\eta^2 = 0.21$<br>interaction: $F(1,38) = 4.37$, $p = \mathbf{0.043}$, partial $\eta^2 = 0.1$ | |
| | ns | post-hoc test $p_{\text{tukey}} = \mathbf{0.003}$<br>Cohen's $d = -1.14$ |
| Improvement in proportion of optimal actions (5B) | age: $F(1,38) = 5.1$, $p = 0.03$, partial $\eta^2 = 0.12$<br>interaction: $F(1,38) = 4$, $p = 0.052$, partial $\eta^2 = 0.095$ | |
| | ns | post-hoc test $p_{\text{tukey}} = \mathbf{0.018}$<br>Cohen's $d = -0.95$ |
| Q-learning-alpha (6A) | age: $F(1,38) = 0.04$, $p = 0.84$, partial $\eta^2 = 0.001$<br>interaction: $F(1,38) = 0.26$, $p = 0.62$, partial $\eta^2 = 0.007$ | |
| Q-learning-tau (6A) | age: $F(1,38) = 1$, $p = 0.32$, partial $\eta^2 = 0.026$<br>interaction: $F(1,38) = 0.034$, $p = 0.86$, partial $\eta^2 = 0.0009$ | |
| Secondary parameters – experiment 2 |  |  |
| Proportion of optimal actions – adjacent (S1B) | age: $F(1,38) = 0.97$, $p = 0.33$, partial $\eta^2 = 0.025$;<br>interaction: $F(1,38) = 1.93$, $p = 0.17$, partial $\eta^2 = 0.05$ | |
| Proportion of optimal actions – distant (S1B) | age: $F(1,38) = 5.32$, $p = \mathbf{0.027}$, partial $\eta^2 = 0.12$;<br>interaction: $F(1,38) = 12.84$, $p < \mathbf{0.001}$, partial $\eta^2 = 0.25$ | |
| | | post-hoc test $p_{\text{tukey}} < \mathbf{0.001}$,<br>Cohen's $d = -1.3$ |
| Improvement in action entropy - distant (S2B) | age: $F(1,38) = 7.73$, $p = \mathbf{0.008}$, partial $\eta^2 = 0.17$<br>interaction: $F(1,38) = 0.21$, $p = 0.65$, partial $\eta^2 = 0.006$ | |
| Memory questionnaire – adjacent (S2) | age: $F(1,38) = 0.03$, $p = 0.86$, partial $\eta^2 = 0.0008$<br>interaction: $F(1,38) = 0.6$, $p = 0.44$, partial $\eta^2 = 0.016$ | |
| Memory questionnaire – distant (S2) | age: $F(1,38) = 110.37$, $p < \mathbf{0.001}$, partial $\eta^2 = 0.74$<br>interaction: $F(1,38) = 0.006$, $p = 0.94$, partial $\eta^2 = 0.0004$ | |

**Table S3.** Summary of all correlation measurements.

| Parameter 1 | Parameter 2 | Short ISI | Long ISI |
| --- | --- | --- | --- |
| Improvement in number of episodes completed | Q-learning alpha (4B) | $r(64) = 0.4$, $p < 0.001$ | ns |
| Improvement in number of episodes completed | Q-learning tau (4B) | $r(64) = -0.4$, $p < 0.001$ | ns |
| Proportion of optimal actions | Q-learning alpha (6B) | ns | ns |
| Proportion of optimal actions | Q-learning tau (6B) | $r(38) = -0.38$, $p = 0.015$ | $r(38) = -0.5$, $p = 0.008$ |
| Improvement in proportion of optimal actions | Improvement in action entropy - distant (S3B) | Not tested | $r(38) = -0.63$, $p < 0.001$ |

